# DALI (Diversity AnaLysis Interface): a novel tool for the integrated analysis of multimodal single cell RNAseq data and immune receptor profiling

**DOI:** 10.1101/2021.12.07.471549

**Authors:** Kevin Verstaen, Kevin Ibech, Inés Lammens, Jana Roels, Yvan Saeys, Bart N Lambrecht, Niels Vandamme, Stijn Vanhee

**Affiliations:** Data Mining and Modeling for Biomedicine, VIB Center for Inflammation Research, Ghent, BE; Department of Applied Mathematics, Computer Science and Statistics, Ghent University, Ghent, BE; Department of Internal Medicine and Pediatrics, Ghent University, BE; Laboratory for Immunoregulation and Mucosal Immunology, VIB Center for Inflammation Research, Ghent, BE; VIB Single Cell Core Facility, BE

**Author notes:** equal contribution (Kevin Verstaen, Kevin Ibech, Niels Vandamme, Stijn Vanhee).

## Abstract

Single-cell RNA sequencing is instrumental to unravel the cellular and transcriptomic heterogeneity of T and B cells in health and disease. Recent technological advances add additional layers of information allowing researchers to simultaneously explore the transcriptomic, surface protein and immune receptor diversity during adaptive immune responses. The increasing data complexity poses a burden on the workload for bioinformaticians, who are often not familiar with the specificities and biology of immune receptor profiling. The wet-lab modalities and sequencing capabilities currently have outpaced bioinformatics solutions, which forms an ever-increasing barrier for many biologists to analyze their datasets. Here, we present DALI (**D**iversity **A**na**L**ysis **I**nterface), a software package to identify and analyze T cell and B cell receptor diversity in high-throughput single-cell sequencing data. DALI aims to support bioinformaticians with a functional toolbox, allowing seamless integration of multimodal scRNAseq and immune receptor profiling data generated through 10X Genomics Cell Ranger software. The R-based package builds further on workflows using the Seurat package and other existing tools for BCR/TCR analyses. In addition, DALI is designed to engage immunologists having limited coding experience with their data, using a browser-based interactive graphical user interface. The implementation of DALI can effectively lead to a two-way communication between wet-lab scientists and bioinformaticians to advance the analysis of complex datasets.

## Introduction

Dissecting heterogeneity of immune cell responses in disease is at the heart of immunological research. With the development of single cell sequencing methods, the ability to distinguish cellular heterogeneity based on RNA expression became a popular tool for immunological research. Recent developments in these methods have added additional layers of information, including surface protein expression (CITE-Seq) and immune receptor profiling (VDJseq). Understanding the evolution of immune receptor profiles in adaptive immune responses is essential to fully appreciate the heterogeneity of adaptive immune responses in the fields of infectiology, allergy, autoimmunity, cancer and vaccinology.

Both B lymphocytes and T lymphocytes express immune receptors with an astonishing diversity, forming an anticipatory repertoire to fend off any antigenic exposure. Throughout their development, diversity is induced in the B cell receptor (BCR) heavy and light chains and the T cell receptor (TCR) alpha and beta chains or gamma and delta chains. During this rearrangement process genomic segments of the variable (V), diversity (D) and joining (J) regions are recombined in a random fashion, which are then combined with constant (C) region sections. In addition, at the junctions of these recombined fragments, random nucleotides are inserted, increasing the diversity even more. The anticipatory repertoire of B and T cells is estimated to contain over 10^13^ unique receptor sequences (1). Upon antigen exposure, B cells will develop into either memory cells or antibody secreting plasma cells. The main goal here is to produce antibodies with very high specificity towards the antigen. This is induced by introducing random mutations into the recombined BCR sequences, increasing antigen affinity in a stochastic manner. To this end, B cells undergo somatic hypermutation (SHM) and subsequent selection for receptor affinity in germinal center reactions. Hence, these processes further increase the complexity of the immune receptor repertoire.

While we have seen rapid technological advances now allowing us to sequence the immune receptor repertoires of hundreds of thousands of cells, the tools that allow easy and meaningful analysis of these datasets are not developing at the same speed. To help the field in accommodating this need, we have developed the Diversity AnaLysis Interface (DALI), which interacts with the Seurat R package (2) and is aimed to support the advanced bioinformatician with a set of novel methods and an easier integration of existing tools for BCR and TCR analysis in their single cell workflow. With this, we aim to produce meaningful output from complex datasets. On the other hand, DALI aids immunologists with a limited background in bioinformatics through an interactive R Shiny app based graphical user interface (GUI), allowing easy data exploration. The implementation of DALI can effectively lead to a two-way communication between wet lab scientists and bioinformaticians, and advance the analysis of complex datasets

## Results

### Supported platforms

DALI can be used on any single-cell dataset processed with the R Seurat package and for which there is corresponding immune profiling data. The package has been built and tested based on the 10x Genomics cellranger vdj and cellranger multi output. However, any platform that generates AIRR compliant output (3) can be integrated into an immune receptor annotated R Seurat object.

### DALI analysis pipeline

DALI can be integrated in an existing single-cell workflow in a couple of different ways, as shown in figure 1. First, an existing R Seurat object containing single cell gene expression data can be linked to its corresponding immune receptor profiling data via the interactive DALI Shiny app (Fig 1A, step A). This can be done by simple selection of the source R Seurat object and the folder containing the immune receptor sequencing output generated by cellranger vdj or cellranger multi (Figure 1B). DALI allows to load both BCR and TCR data in the same object and allows for seamless switching between the 2 data modalities. Secondly, an extended R Seurat object can be generated outside the interactive tool with the provided tools in the DALI package, in which the immune receptor data is linked to its corresponding cell (Figure 1A, step B). This extended R Seurat object can then be loaded in the shiny application (Figure 1A, step B’) or be used in the R environment for further analysis using DALI or any other R tool. With the increasing amount of single cell sequencing datasets that are being generated, we have also accommodated integration of immune receptor data with their respective gene expression data in the case of multiple samples. Alternatively, the DALI interactive GUI could also be used without any immune receptor data. This allows biologists to explore the gene expression data in an interactive way.

**Figure 1:**
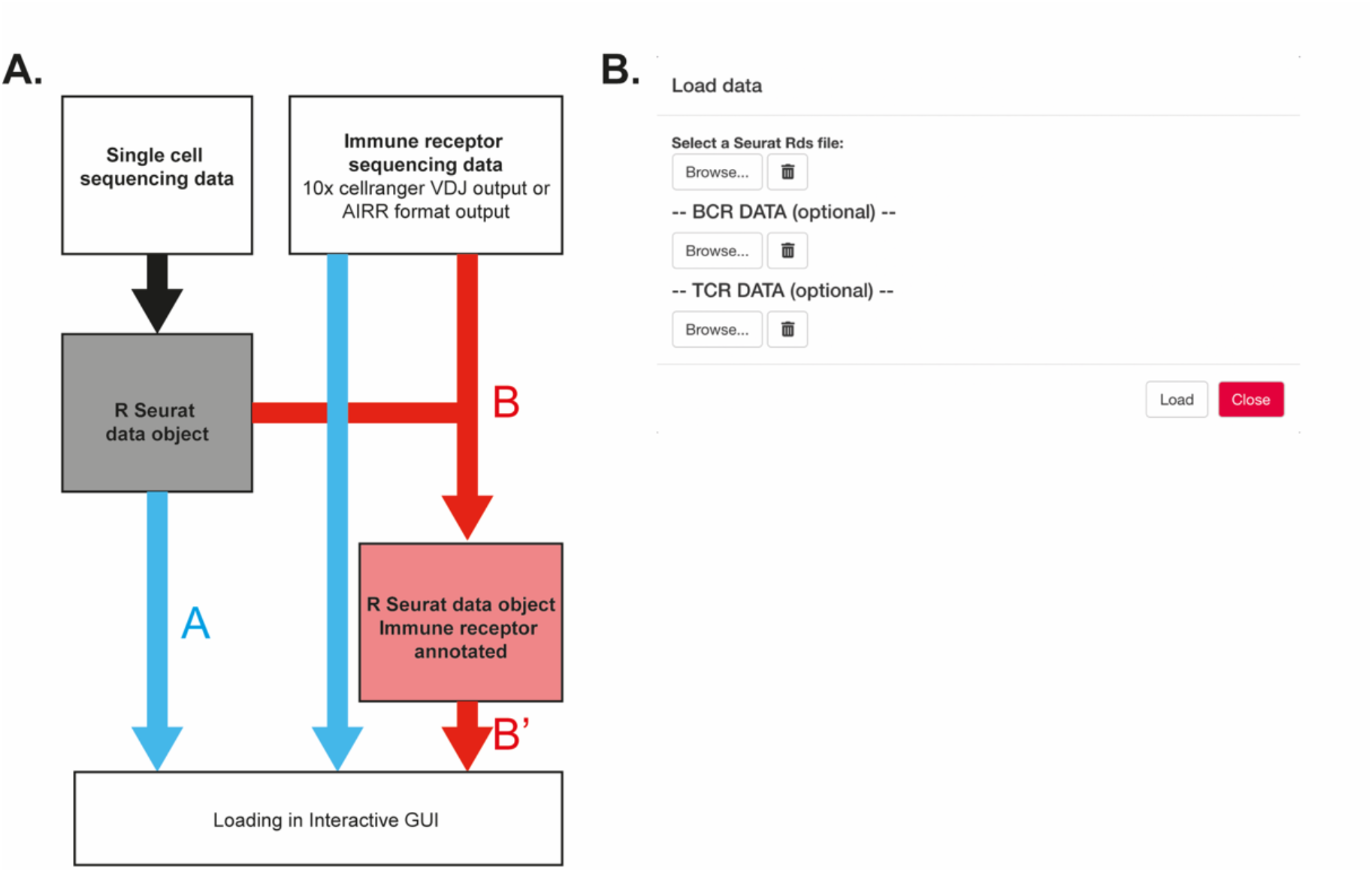
Overview of DALI pipelines using either **(A)** loading of the R Seurat object and immune profiling data directly into the interactive Shiny app **(B)** an extended R Seurat object containing immune receptor profiling data, which can be loaded into the interactive Shiny app (B’).

### Overview of the Shiny application

Most of the functionalities of DALI are bundled in a user-friendly Shiny application, which can be run in the browser. Here, the user has the option to go through different steps adding up to an in-depth analysis of the integrated (multimodal) single cell dataset. The user can navigate through the **General View** to view immune receptor gene segment usage plotted on the dimensionality reductions available in the object (Figure 2A). Next, the **Clone View** allows for an instantaneous assessment of clonotype or clone expansion and potential clonal overlap between clusters using the clonotype connection plots (Figure 2B). Additionally, in-depth information on the top clonotypes within each of the metadata factors can be obtained. Next, using the **Population comparison view**, users can directly compare immune receptor properties between patient, condition, treatment or any other metadata variable available (Figure 2C). Once a clonotype of interest is identified, the **Clonotype view** tab can be used to obtain full length information on these clonotypes (Figure 2D). This is of great interest to generate recombinant monoclonal antibodies, perform receptor modeling or to generate immune receptor transgenic cells/animals. The clonotype view allows for interactive generation of hierarchical clusters of BCRs, denoting ongoing somatic hypermutation of one clonotype. The nodes can be colored based on gene expression data or any meta data parameter of interest. In addition, the clonotype view allows for visualization of somatic hypermutation. In the **Transcriptomics view**, expression of the genes of interest can be plotted on the dimensionality reduction plots and linked to the position of the clonotype of interest (Figure 2E). In the **Differential Gene Expression view**, differential gene expression can be calculated between cells using any metadata parameter, including immune receptor properties (Figure 2F). The results of this analysis are visualized in a volcano plot to quickly identify up- or downregulated genes.

**Figure 2:**
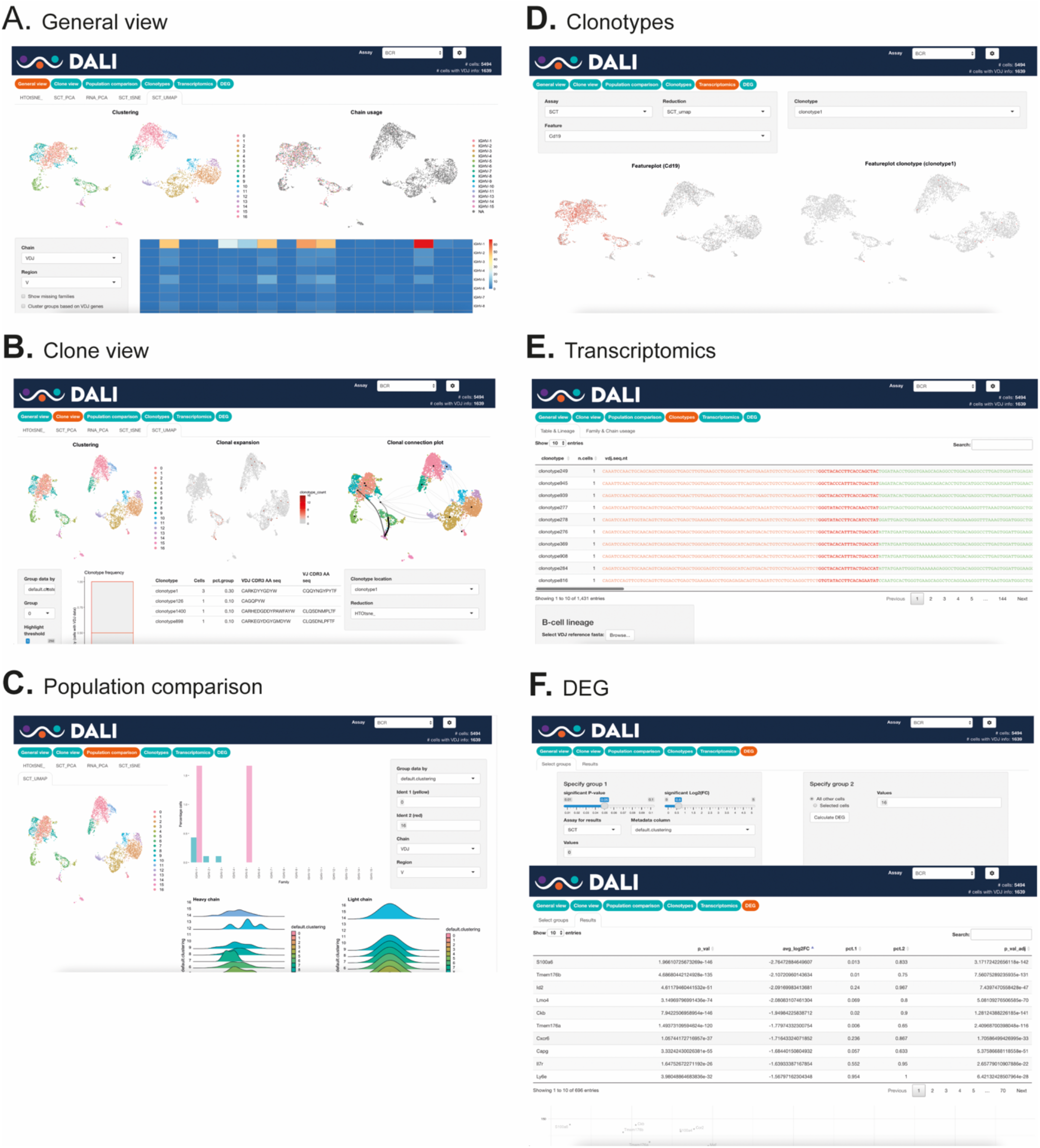
Overview of the different DALI Shiny app tabs. **(A)** General View, **(B)** Clone View, **(C)** Population comparison, **(D)** Clonotype view, **(E)** Transcriptomics view, **(F)** Differential Gene Expression view.

### Advanced DALI functionality

#### Trajectory analysis

The DALI package has two functions to perform trajectory analysis on a given dataset. Making use of the slingshot package (4), the trajectory can either be rooted on a specified cluster or a cluster with the highest expression of a specified feature. Alternatively, Dynverse (5) can be used to unlock a wider variety of trajectory inference methods.

### A test case: analysis of house dust mite driven B cell responses in mediastinal lymph nodes

As a test case, we have performed single cell immune profiling of both B and T cell receptors of enriched subsets of B and T cells from mediastinal lymph node. These cells were isolated from mice sensitized to house dust mite extract (HDM) and subsequently challenged with HDM for 3 weeks. During the adaptive immune response in the lymph node in response to allergen exposure, a germinal center response will be initiated (Figure 3). During this process, naïve B cells and T cells will be activated and will differentiate into germinal center B cells (GCB) and follicular helper T cells (TFH), respectively. Germinal center B cells will increase the affinity of their B cell receptors and undergo receptor class switching in the dark zone of the germinal center. Upon exit of the dark zone, B cells enter the germinal center light zone where their newly obtained receptor affinity is tested. If successful in competing for limiting amounts of antigen, the light zone B cell will enter the dark zone again for another round of affinity maturation. From these germinal center responses, dedicated antibody producing cells (plasma cells) or memory B cells can arise, the latter effectively forming a memory of previous antigen exposure. We will here define dark zone B cells (DZB) and light zone B cells (LZB), memory B cells (Bmem), as well as plasma cells (PC). In addition, we zoom in at clonal evolution of a clone of interest and we will define antigen specific B cells and analyze their immune receptor repertoires.

**Figure 3:**
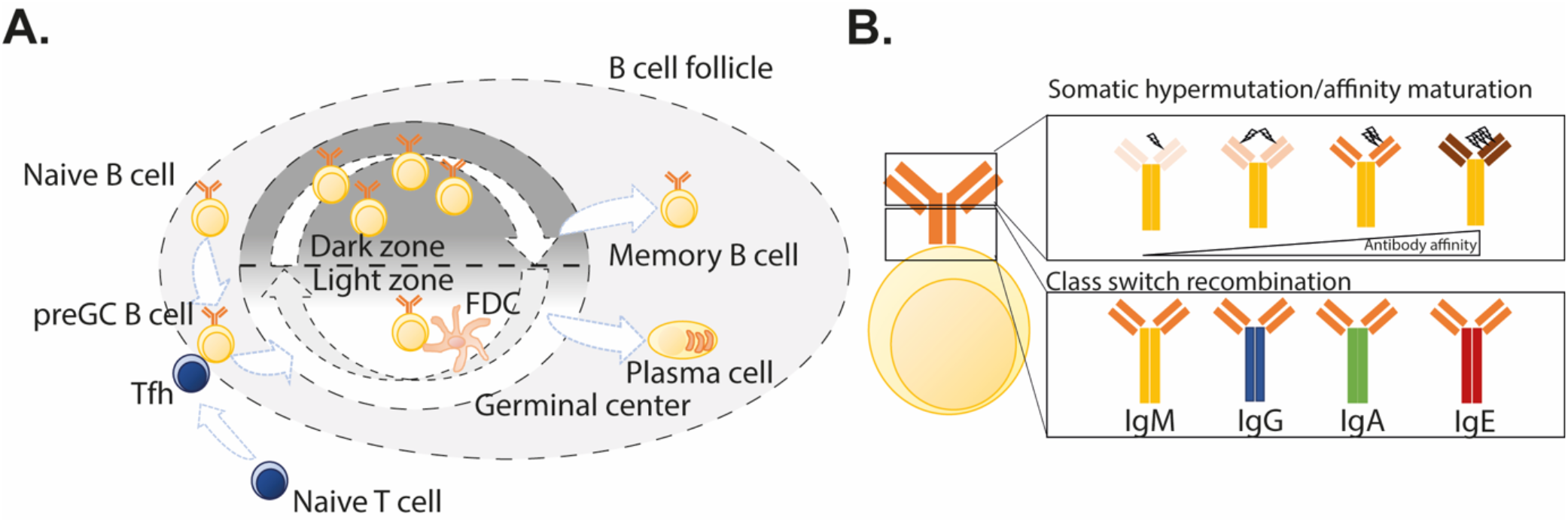
Overview of B cell responses **(A)** B cell germinal center (GC) responses. An activated B cell will interact with its cognate follicular helper T cell (Tfh). Upon this interaction, the B cell will differentiate into a germinal center B cell (GCB) and migrate into the GC dark zone. Here, the B cell will proliferate and can undergo class switch recombination and somatic hypermutation directed by the AID protein. The edited receptor is then tested for its potential to bind antigen presented by follicular dendritic cells (FDC). The GCB can undergo multiple iterations of the cycle. From the GC response, memory B cells and plasma cells can arise. The former will quickly respond and re-enter a GC reaction upon future antigenic exposure, while plasma cells are dedicated antibody producing cells. **(B)** Schematic overview of B cell receptor (BCR) affinity maturation and class switch recombination. With increasing somatic hypermutation, the binding affinity of the BCR/secreted antibody might be increased. Upon class switch recombination, the BCR/antibody constant region is switched, providing the receptor with different signaling capacities and the secreted antibody with different functional properties.

#### Definition of DZB, LZB and PC

We set out to define different B cell subsets using the DALI interactive mode (Figure 4A), we defined B cells through Cd79a expression through the transcriptomics view (Figure 4B). To define germinal center B cells, we defined cells expressing Bcl6 and Mki67 (Figure 4C). These cells can be traced back to cluster 7 (Figure 4A). We defined LZB within cluster 7 by CD86 and Fcer2a expression (Figure 4D), while DZB were defined using Hmmr and Aicda expression (Figure 4E). Plasma cells were defined by Xbp1 and Jchain expression (Figure 4F). Lastly, Bmem were defined using high expression of Klf2 and Bach2 (Figure 4G). Using the general view, we can next determine heavy and light chain constant region usage, where we see evidence for class switching towards IgG1 and IgG2b in cluster 7 (Figure 4H). Using the clone view, we can see that there is clonal expansion within cluster 7 (Figure 4I) and that some clones are shared between cluster 7, cluster 0 and cluster 15 (Figure 4J).

**Figure 4:**
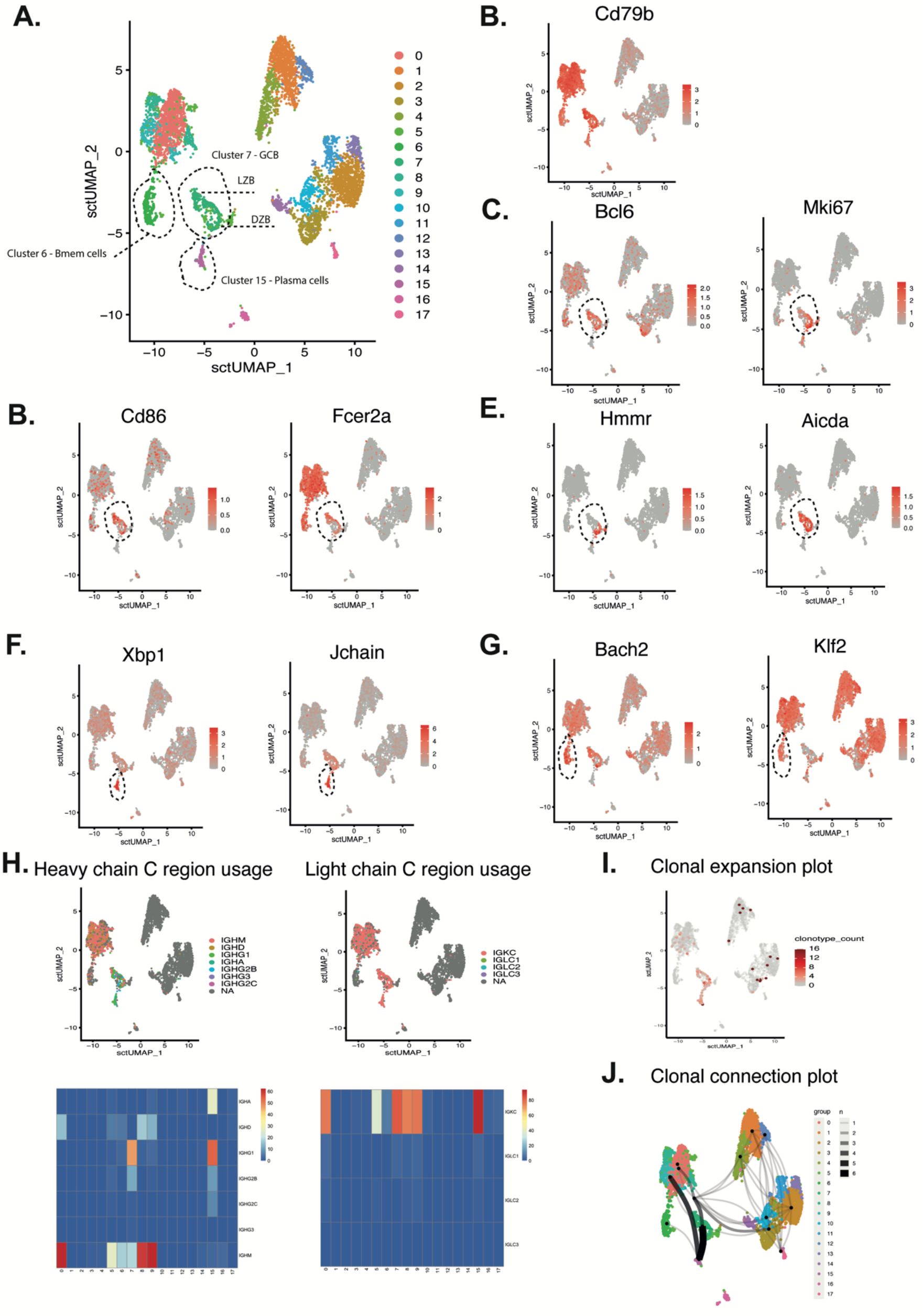
Definition of B cell subsets using DALI. **(A)** Definition of the cluster containing different B cell subsets. **(B)** Definition of CD79a expression B cells **(C)** Definition of germinal centers B cells by Aicda and Mki67. **(D)** Definition of light zone B cells (LZB) by CD86 and Fcer2a expression. **(E)** Definition of dark zone B cells (DZB) by Hmmr and Aicda expression. **(F)** Definition of plasma cells by Xbp1 and Jchain expression. All populations were defined using the transcriptomics view. **(G)** Definition of Bmem by Bach2 and Klf2 expression. All populations were defined using the DALI transcriptomics view **(H)** DimPlots and heatmaps depicting B cell receptor heavy and light chain constant region usage from the DALI general view, **(I)** Clonotype expansion plot from the clone view, **(J)** Clonotype connection plot from the DALI clone view.

#### Understanding clonal evolution

Next, we aim to understand the clonal evolution of some of the expanded clones in cluster 7. To this end, we define an expanded clone in the clone view, and define this clone further in the clonotype view. Here we zoom in on clonotype 2 (Figure 5A, B). Using our tree analysis in the clonotype view, we can appreciate the clonal evolution of clonotype 2 (Figure 5C,D). Here we highlight the evolution of class switching within this tree view (Figure 5C) and the evolution of AID expression (Figure 5D).

**Figure 5:**
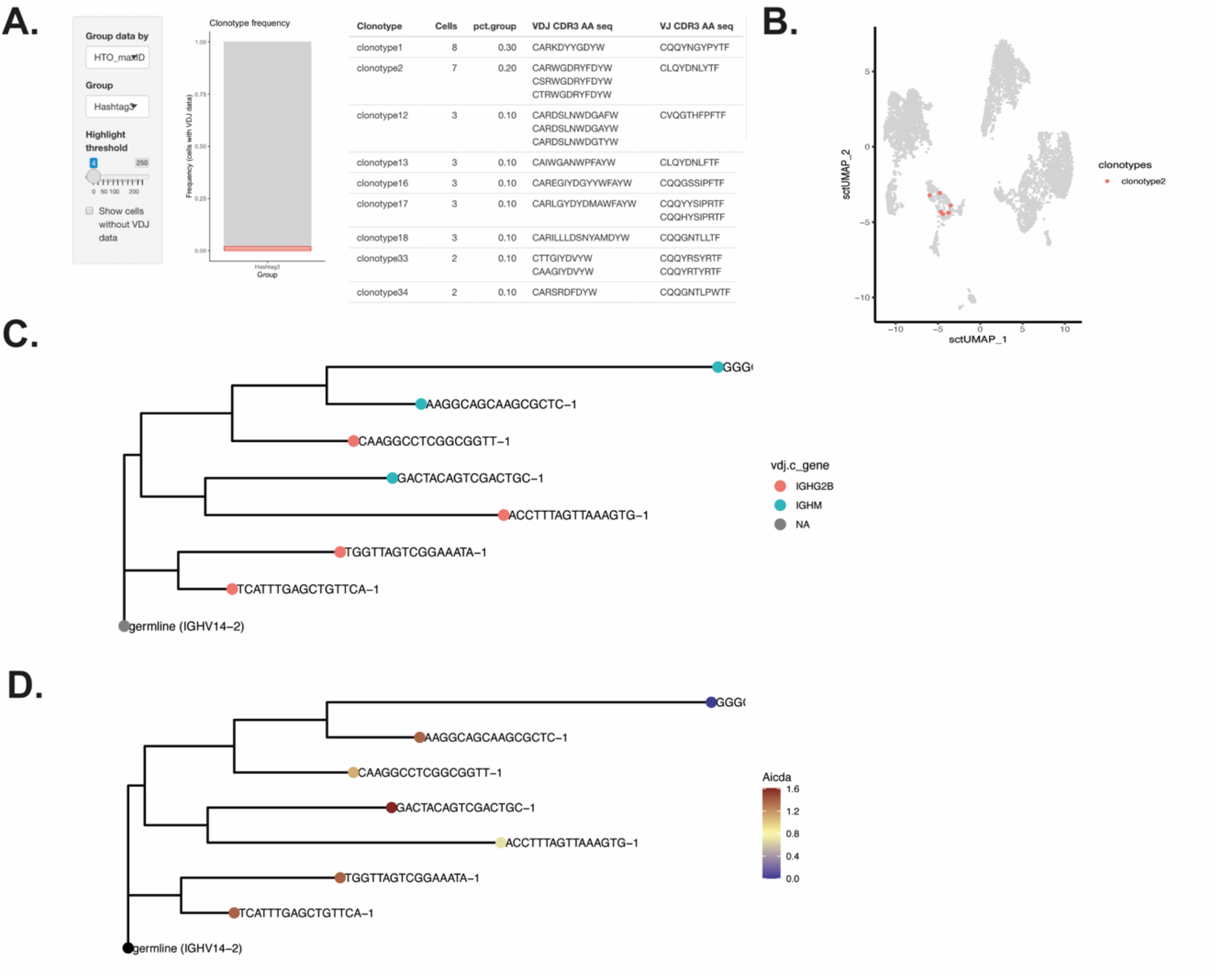
Analysis of clonal evolution. **(A)** Definition of expanded clones using the clone view, **(B)** Localization of selected clones on the UMAP plot (clonotype2 is highlighted), **(C)** Hierarchical clustering of clonotype2, with node ends highlighting the isotype class, **(D)** Hierarchical clustering of clonotype2, with node ends highlighting the Aicda expression levels.

#### Analysis of antigen specific B cells

Finally, we analyze the repertoire of Der p2 specific B cells, a major allergen in house dust mite. Using FACS, we sorted Der p2 reactive cells using Der p2 protein tetramers (Figure 6A), and upon sample hashing, combined these for single cell sequencing. Demultiplexing of the samples shows the location of the two Der p2+ samples on the UMAP (Figure 6B). Using the clone view, we can obtain information on the Der p2 reactive BCR sequences (Figure 6C) and obtain detailed information on the clonotype in the clonotype view (Figure 6D).

**Figure 6:**
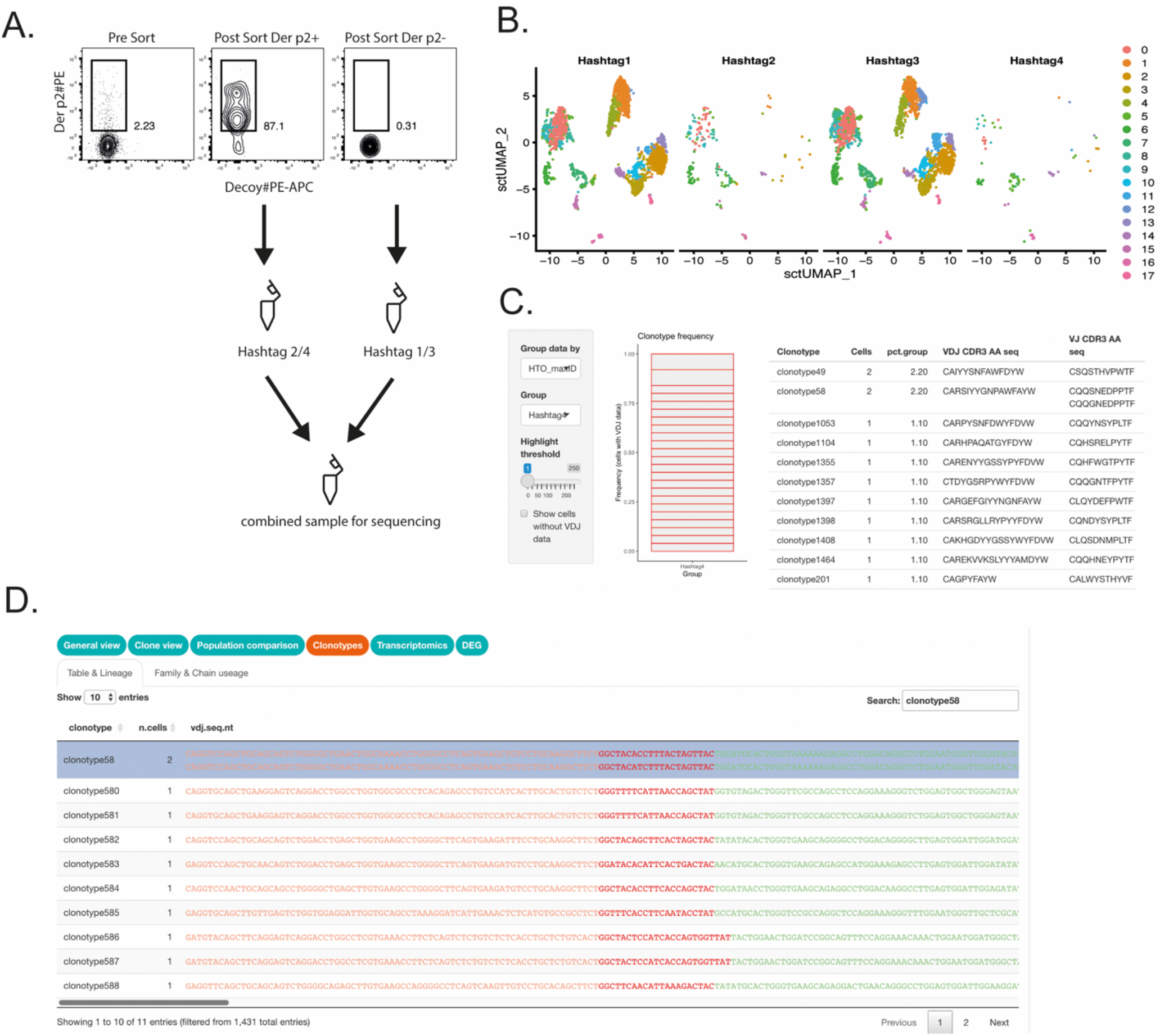
Analysis of antigen reactive B cells. **(A)** Isolation of Der p2 reactive B cells from mediastinal lymph node by FACS using a Der p2 protein tetramer (FACS plots show CD19+ pregated cells), the sample hashing strategy is shown **(B)** Demultiplexing of the sample based on hashtag expression, **(C)** Definition of Der p2 reactive clones in sample 4, **(D)** Detailed information on one of the clonotypes shown in C.

### Conclusion and future direction

We here provide a comprehensive R based tool for the integrated analysis of single cell gene expression data and immune receptor profiling in the Seurat ecosystem. One of the unique aspects of this tool is both the availability of a toolset in the R environment for the experienced bioinformatician, as well as a GUI based interface for the biologist with limited bioinformatics background. While we strongly believe that this will aid the field and reduce the generation of data black boxes, we do realize that no tool is complete, and that the GUI analysis might be limited to obtain complete understanding of the dataset. Nonetheless we believe that this way of analyzing data provides a first step, upon which the biologist can interact with experienced bioinformaticians. DALI is also fully open source, which allows the community to aid in the further development of this tool. Lastly, DALI will receive regular updates to accommodate the rapid evolutions in the field of single cell transcriptomics.

## Methods

### Scripting and computational analysis

All scripting was performed using R (6). All analyses were performed using R version 4.2.0 “Vigorous Calisthenics” and DALI version 2.0.0.

### Mice and treatments

All mice were obtained through Janvier (France) and were of the C57BL6 strain. Mice were sensitized with 1ug house dust mite extract (ALK, Denmark) in 80ul of PBS instilled intratracheally and challenged 3 times a week for 3 weeks with 50ug of house dust mite extract in 40ul PBS intranasally. These methods are described in greater detail in (7).

### Single cell sequencing

#### Single-cell RNA and BCR/TCR sequencing experiments

Upon isolation of mediastinal lymph nodes from 2 biological replicates, cells were stained with CD19 Pe-Cy5 (clone 1D3), Fixable viability dye Live/Dead (L/D) Efluor 506, homemade Der p2 protein tetramers coupled to streptavidin-PE and biotin-streptavidin-PE-AF647 decoys. This staining was performed on ice for 30’ in PBS containing 3% of FCS. After sorting Der p2 reactive and non-reactive cells, the cells were stained with TotalSeq-C hashing antibodies (BioLegend) diluted 1:500. 4 samples were multiplexed per lane using TotalSeq-C Cell Hashing Antibodies. Sorted cells were mixed according to the following ratio: non-B cells/B cells/Der p2 reactive B cells in a 45:45:10 ratio. Samples from 2 biological replicates were then mixed in a 1:1 ratio. Sorted single-cell suspensions were resuspended at an estimated final concentration of 2000 cells/μl and loaded on a Chromium GemCode Single Cell Instrument (10x Genomics) to generate single-cell gel beads-in-emulsion (GEM). The scRNA/Feature Barcoding/BCR/TCR libraries were prepared using the GemCode Single Cell 5’ Gel Bead and Library kit, version 1.1 (10x Genomics, Cat. 1000165) according to the manufacturer’s instructions. The cDNA content of pre-fragmentation and post-sample index PCR samples was analyzed using the 2100 BioAnalyzer (Agilent). Sequencing libraries were loaded on an Illumina NovaSeq flow cell at VIB Nucleomics core with sequencing settings according to the recommendations of 10x Genomics, pooled in a 75:20:5 ratio for the gene expression, TCR/BCR and antibody-derived libraries, respectively.

#### Single-cell data processing and analysis

The Cell Ranger pipeline (10x Genomics, version 6.1.1) was used to perform sample demultiplexing and to generate FASTQ files for read 1, read 2 and the i7 sample index for the gene expression and cell surface protein libraries. Read 2 of the gene expression libraries was mapped to the mouse reference genome (GRCm38; Ensembl release 99). Subsequent barcode processing, unique molecular identifiers filtering, and gene counting was performed using the Cell Ranger suite version 6.1.1. Outlier cells were identified based on 3 metrics (library size, number of expressed genes and mitochondrial proportion) and cells were tagged as outliers when they were 3 median absolute deviation (MADs) away from the median value of each metric across all cells. Low quality cells (low UMI counts, high % of mitochondrial genes) were removed from the analysis.

## Data availability

The presented dataset is available through our Github page https://github.com/vibscc/DALI

## Acknowledgements

We would like to thank Dr Nincy Debeuf, Dr. Andrew Brown and Sam Dupont for testing beta versions of DALI.

We are grateful for the excellent support received from the VIB Flow Core Facility, VIB Nucleomics Core Facility, VIB Protein Core Facility and the VIB Single Cell Core Facility. https://vib.be/core-facilities

SV is supported by a Fonds voor Wetenschappelijk Onderzoek Vlaanderen (FWO) senior postdoctoral fellowship FWO-1244321N

## Author contributions

KV conceived the project, wrote the code and wrote the manuscript, KI wrote the code, IL performed experiments, JR wrote the code, YS, BNL, NV conceived the project and wrote the manuscript, SV conceived the project, wrote the code, wrote the manuscript, performed experiments, analyzed data and supervised the study.

